# Short superficial white matter and aging: a longitudinal multi-site study of 1,293 subjects and 2,711 sessions

**DOI:** 10.1101/2022.06.06.494720

**Authors:** Kurt G Schilling, Derek Archer, Fang-Cheng Yeh, Francois Rheault, Leon Y Cai, Andrea Shafer, Susan M. Resnick, Timothy Hohman, Angela Jefferson, Adam W Anderson, Hakmook Kang, Bennett A Landman

## Abstract

It is estimated that short association fibers, or “U-shaped” fibers running immediately beneath the cortex, may make up as much as 60% of the total white matter volume. However, these have been understudied relative to the long-range association, projection, and commissural fibers of the brain. This is largely because of limitations of diffusion MRI fiber tractography, which is the primary methodology used to non-invasively study the white matter connections. Inspired by recent anatomical considerations and methodological improvements in U-fiber tractography, we aim to characterize changes in these fiber systems in cognitively normal aging, which provide insight into the biological foundation of age-related cognitive changes, and a better understanding of how age-related pathology differs from healthy aging. To do this, we used three large, longitudinal and cross-sectional datasets (N = 1293 subjects, 2711 sessions) to quantify microstructural features and length/volume features of several U-fiber systems. We find that axial, radial, and mean diffusivities show positive associations with age, while fractional anisotropy has negative associations with age in superficial white matter throughout the entire brain. These associations were most pronounced in the frontal, temporal, and temporoparietal regions. Moreover, measures of U-fiber volume and length decrease with age in a heterogenous manner across the brain, with prominent effects observed for pre- and post-central gyri. These features, and their variations with age, provide the background for characterizing normal aging, and, in combination with larger association pathways and gray matter microstructural features, may provide insight into fundamental mechanisms associated with aging and cognition.

## Introduction

Superficial white matter (SWM) is the layer of white matter just beneath the cortex, and is composed of short association U-shaped fibers, or U-fibers, that primarily connect adjacent gyri. These U-fibers represent a majority of the connections of the human brain [1, 2], occupy as much as 60% of the total white matter volume [1], are among the last parts of the brain to myelinate [3], and contain a comparatively high density of interstitial white matter neurons relative to other white matter[4]. The SWM serves a critical role in brain function [5], plasticity, development, and aging, and is especially affected in disorders such as Alzheimer’s disease [6, 7], autism [8], and schizophrenia [9].

Despite its prevalence and significance, SWM has been understudied relative to the long-range association, projection, and commissural fibers of the brain. This is largely because of the limitations of diffusion MRI fiber tractography [10–12], which is the primary methodology used to non-invasively study the white matter connections [13]. The study of U-fibers using tractography faces anatomical and methodological challenges including partial volume effects, complex local anatomy, and a lack of consensus on definition and taxonomy [12], which complicate development and validation of algorithms dedicated to studying these fiber systems. However, recent innovation in diffusion MRI imaging, processing, and tractography methodologies [10, 12, 14–16] have made it possible to reliably study SWM in health and disease [9, 17–21].

One promising avenue of exploration is to study U-fibers during aging. Studies of the aging brain may provide insight into the biological foundation of age-related cognitive changes, and a better understanding of how abnormal aging (e.g., age-related neurodegenerative disorders) differs from healthy aging [22]. A large body of magnetic resonance imaging (MRI) research has shown that the structure of the human brain is constantly changing with age. In the gray matter, structural MRI studies have shown heterogenous patterns of normal age-related changes in cortical volume and thickness [23–30], with detectable differences in abnormal aging and disease [30–35]. In the white matter, diffusion tensor imaging (DTI) analysis has shown that fractional anisotropy (FA) is negatively associated with age and mean diffusivity (MD) is positively associated with age across several white matter pathways [36–39], and tractography analysis has shown that the volume and surface areas of many pathways decreases with age [40]. These findings have been attributed to myelin loss and/or decreased axonal densities and volumes. However, with few exceptions [41–44], studies of white matter brain aging have focused on the deep white matter and larger long-range pathways of the brain.

Inspired by recent anatomical considerations and methodological improvements in U-fiber tractography [12], and lack of studies of SWM during aging, we sought to characterize changes in these fiber systems during normal aging. To do this, we leveraged three well-established cohorts of aging, including two longitudinal cohorts [Baltimore Longitudinal Study of Aging (BLSA) [45], Vanderbilt Memory & Aging Project (VMAP) [46]], and one cross-sectional cohort [Cambridge Centre for Ageing and Neuroscience (Cam-CAN) [47]]. Within these cohorts, we performed automatic tractography segmentation in 82 U-fiber bundles, characterizing both microstructural features and macrostructural features of these SWM systems, to describe associations between these features and age.

## Methods

### Data

This study used data from three datasets, summarized in **Table 1**, and contained a total of 1293 participants (2711 sessions) aged 50-98 years. All datasets were filtered to exclude participants with diagnoses of mild cognitive impairment, Alzheimer’s disease, or dementia at baseline, or if they developed these conditions during the follow-up interval. Finally, datasets were filtered to focus on participants aged 50+, due to limited samples sizes below 50 years old in each dataset.

**Table 1.** This study used 3 longitudinal and cross-sectional datasets, with a total of 1293 participants ( 2711 sessions), aged 50-98 years. Distributions of age at baseline, and number of sessions, are shown for each individual dataset.

| Dataset | Number of Subjects | Number of Sessions | Age |
| --- | --- | --- | --- |
| Baltimore Longitudinal Study of Aging | 741<br>328 M | 1788<br>Range [1 8] | [50 98]<br>74.1 +/- 9.9 |
| Cambridge Centre for Ageing Neuroscience | 365<br>186 M | 365<br>Range [1] | [50 88]<br>68.0 +/- 10.3 |
| Vanderbilt Memory & Aging Project | 187<br>113 M | 558<br>Range [1 4] | [60 95]<br>74.2 +/- 7.0 |
|  | **1293**<br>**627 M** | **2711**<br>**Range [1 8]** | **[50 98]**<br>**73.5 +/- 9.3** |

First, was the Baltimore Longitudinal Study of Aging (BLSA) dataset, with 741 participants scanned multiple times ranging from 1 to 8 sessions, and time between scans ranging from 1 to 10 years, yielding a total of 1788 diffusion sessions. Diffusion MRI data was acquired on a 3T Philips Achieva scanner (32 gradient directions, b-value=700s/mm2, TR/TE=7454/75ms, reconstructed voxel size=0.81×0.81×2.2mm, reconstruction matrix=320×320, acquisition matrix=115× 115, field of view=260×260mm). Second, was data from the Vanderbilt Memory & Aging Project (VMAP), with 187 participants, scanned between 1-4 sessions, with a total of 558 diffusion datasets. Diffusion MRI data was acquired on a 3T Philips Achieva scanner (32 gradient directions, b-value=1000s/mm2, reconstructed voxel size=2×2×2mm). Third, was data from the Cambridge Centre for Ageing and Neuroscience (Cam-CAN) data repository [47] with 356 participants, each scanned once using a 3T Siemens TIM Trio scanner with a 32-channel head coil (30 directions at b-value=1000s/mm2, 30 directions at b-value=2000s/mm2, reconstructed voxel size=2×2×2mm). All human datasets from Vanderbilt University were acquired after informed consent under supervision of the appropriate Institutional Review Board. This study accessed only de-identified patient information.

### Tractography and U-fiber bundle dissection

For every subject and every session, sets of U-fiber pathways were virtually dissected using methodology similar to [12], with small modifications. **Figure 1** visualizes the methodological pipeline. This pipeline utilized MRtrix [48], with tractography performed using the second-order integration probabilistic algorithm [49] to generate 2 million streamlines with a maximum length of 50mm, utilizing anatomical constraints to ensure gray matter to gray matter connections. This pipeline has been shown to result in dense systems of fibers immediately adjacent to the cortical sheet [12].

**Figure 1.**
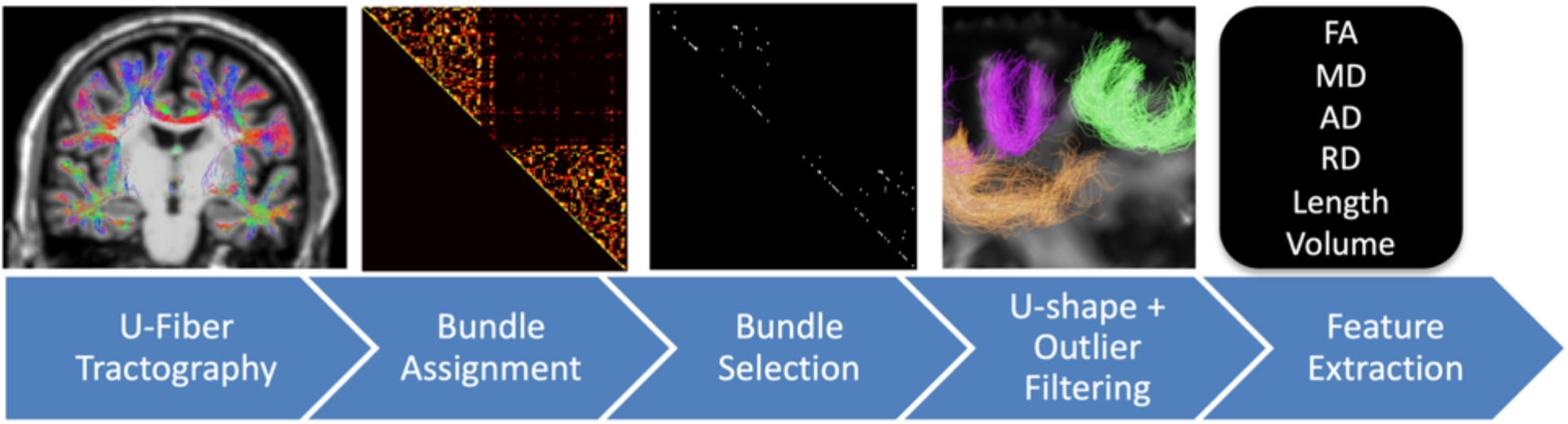
Methodological pipeline. Fiber tractography is constrained based on anatomy and length, and streamlines are assigned to edges in a connection matrix. Only bundles reproducible across the studied population (N=82) are kept for analysis. Bundles are then filtered based on shape and outlier removals. Finally, for each bundle and each subject, microstructural and macrostructural features are extracted for analysis.

Freesurfer [50] was run on the T1-weighted images, and results transformed to diffusion MRI space with ANTs. For this work, we chose to use the Destrieux atlas [51] parcellation, utilizing only the neocortex labels, to assign all streamlines to edges in a connection matrix, resulting in a potential 164×164 SWM bundles. An empirical decision was made to select only those bundles that are reproducible across 75% of the studied population (containing a minimum of 200 streamlines), resulting in 82 U-fiber bundles studied. These bundles were filtered to remove streamlines that were not U-shaped using the scilpy toolbox (https://github.com/scilus/scilpy), and further filtered to remove outlier streamlines [52].

A list of the 82 bundles, using nomenclature derived from the Destrieux atlas, is given in the appendix.

### Feature extraction

From the final 82 bundles for each subject, 6 features were extracted including four DTI microstructural measures of fractional anisotropy (FA), and mean, radial, and axial diffusivities (MD, RD, AD) and two macrostructural measures of length and volume, following the procedures in [53].

### Analytical Plan

To investigate the relationship between age and each WM feature, linear mixed effects modeling was performed, with each (z-normalized) feature, Y, modeled as a linear function of age, *y* = *β*_0_ + *β*_1_*Age* + *β*_2_*Sex* + *β*_3_*TICV* + *β*_3_(1 + *AGE* | *DATASET*) + *β*_4_(*SUB*), where subjects (SUB) were entered as a random effect (i.e., subject-specific random intercept), and subject sex (Sex) and total intracranial volume (TICV) as a fixed effects. Additionally, we modelled the association between age and outcome variable as dataset (DATASET) specific due to expected differences in MR protocols [54–58], and included a dataset specific random slope and intercept. We note that the TICV utilized was calculated from the T1-weighted image from the baseline scan.

Due to multiple comparisons, all statistical tests were controlled by the false discovery rate at 0.05 to determine significance. Results are presented as the beta coefficient of estimate *‘B_1_’*, or in other words “the association of the feature ‘y’ with Age”, which (due to normalization) represents the standard deviation change in feature per year. These measures are derived for each pathway and each feature. Additionally, results may be shown as a percent change per year, derived from the slope normalized by the average value across the aging population (from 50-98), and multiplied by 100, which represents the percent change in feature per year. These measures are derived for each pathway and each feature.

## Results

### U-fiber systems

Example U-fiber systems that were consistently identified across the population are shown in **Figure 2** for a single example subject. In the coronal and axial slices, these fibers run immediately below and adjacent to the cortex in locations and geometries expected traditionally assigned to SWM. In the 3D visualization, U-fibers are represented along a large portion of the gray matter surface. Notably, many U-fiber systems start and end within the same cortical label, which still meets our definition of superficial systems.

**Figure 2.**
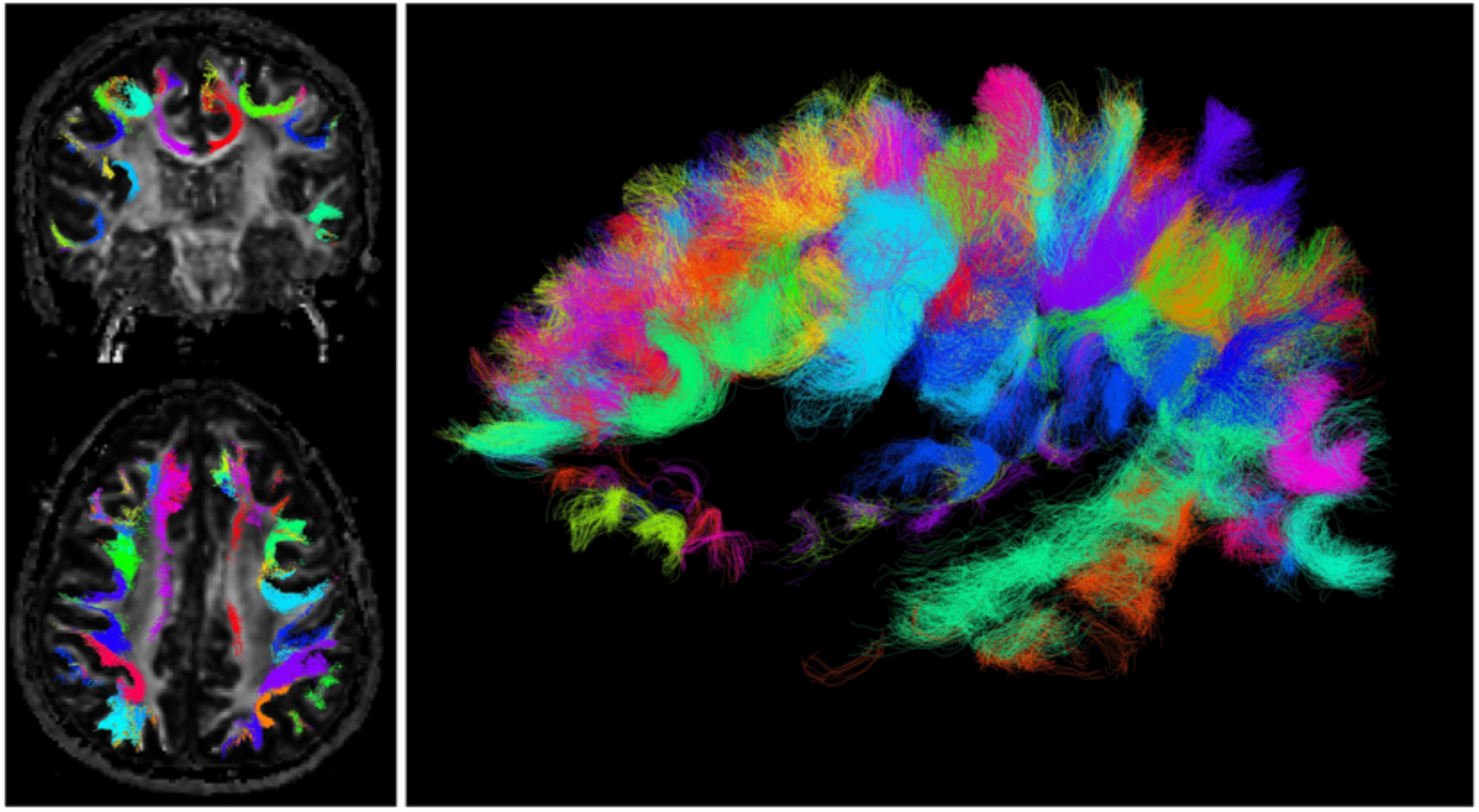
U-fiber systems show expected shape and locations, and cover a large portion of the surface of the brain. 82 U-fibers determined to be robust across a population are shown in a single subject, with distinct colors for each bundle.

### What changes and where?

**Figure 3** shows associations with age of all measures for 7 randomly selected pathways. In line with previous literature in both long association pathways and SWM, FA shows negative associations with age, while the diffusivities show positive associations with age. In general, SWM length and volume tend to decrease with increasing age, even when accounting for TICV, although the effects are not statistically significant for all pathways. As expected, different datasets, with different acquisitions, result in different calculated DTI indices, with much smaller differences in bundle length and volume.

**Figure 3.**
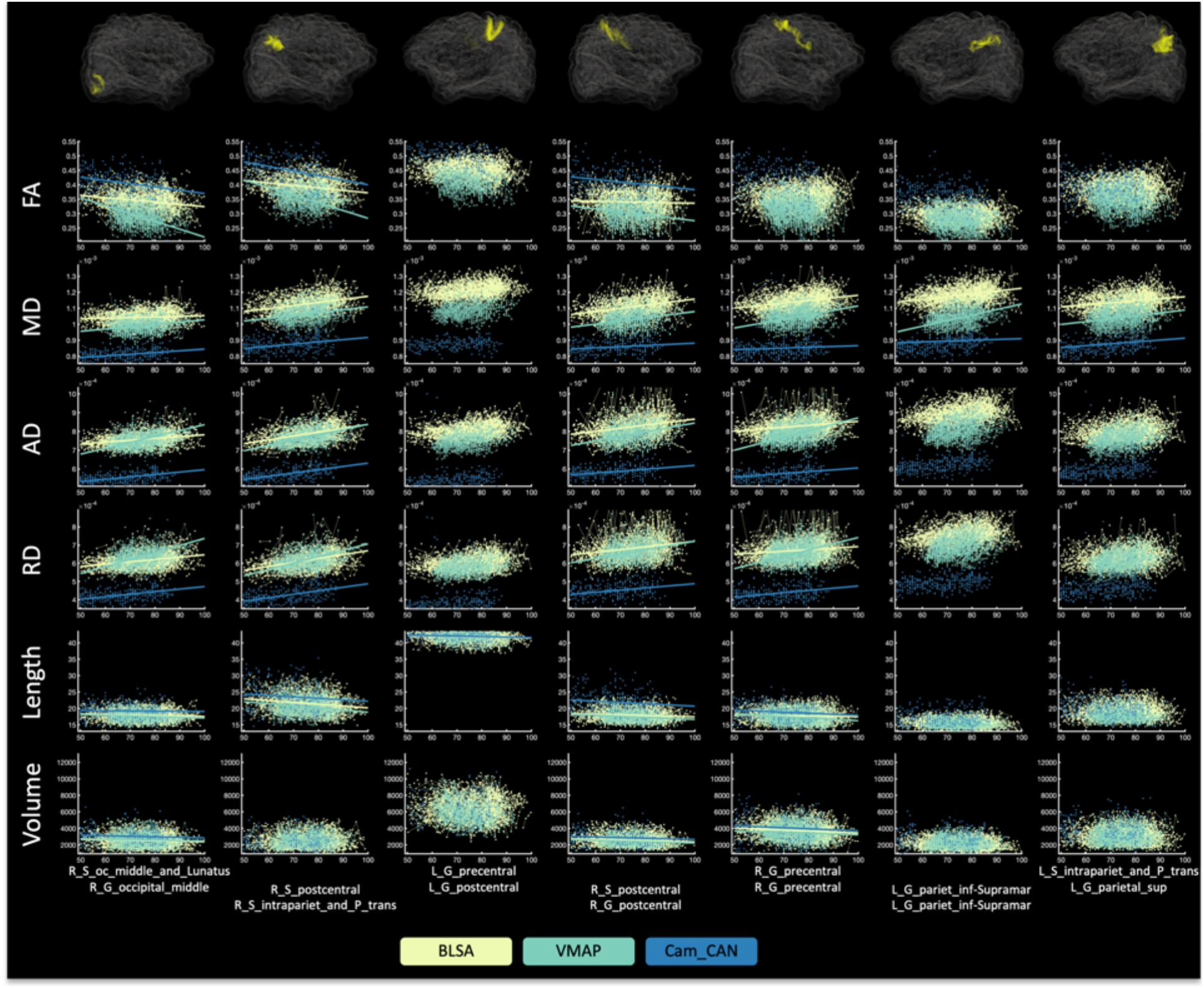
Microstructural and macrostructural features change with age in many pathways. Shown are all studied features for 7 randomly selected pathways, where all data points are shown (with lines connecting longitudinal datasets). A line of best fit is shown if there are statistically significant associations with age, where color indicates the cohort. Visualization of the U-fiber pathways for a single subject are shown overlaid on a transparent brain.

To summarize association with age for all features and all pathways, we show the beta coefficient associations with age for all features in a matrix in **Figure 4**, along with boxplots summarizing the beta coefficients across all studied pathways in **Figure 5**. DTI measures show large, robust associations with age for many pathways. FA in SWM shows negative associations with age, while all diffusivities (AD, MD, RD) show strong positive associations with age. Measures of length and volume show reduced associations with age, for fewer pathways. In general, both length and volume decrease with age for those pathways with statistically significant age associations.

**Figure 4.**
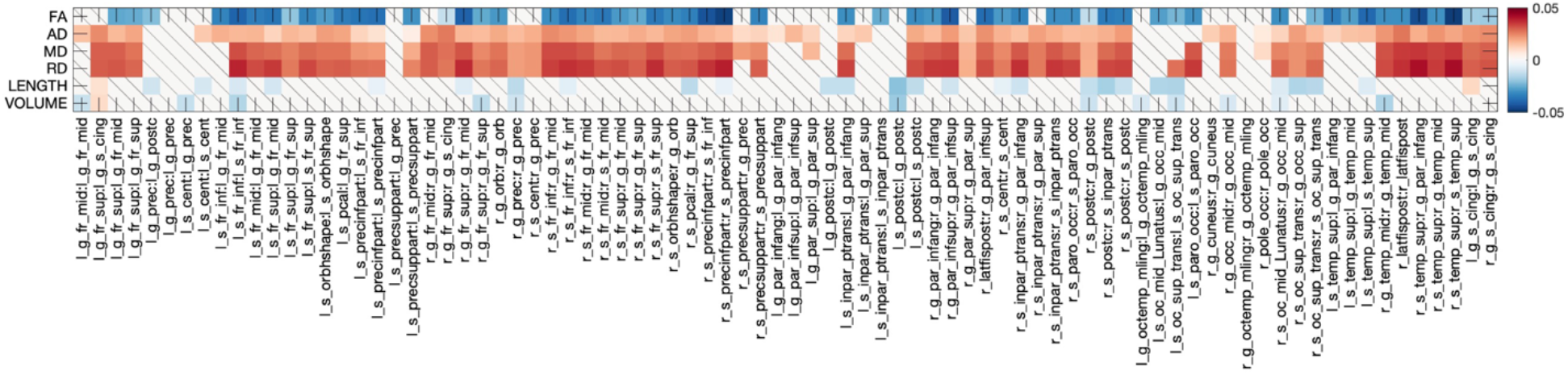
What and where changes occur in SWM during aging. The beta coefficient from linear mixed effects modeling is shown as a matrix for all features across all pathways. Note that the beta coefficient describes *“the association of the feature ‘y’ with Age”, which (due to normalization) represents the standard deviation change in feature per year.* Only those features/pathways with statistically significant age-related changes are colored; non-significant effects are shown as diagonal line.

**Figure 5.**
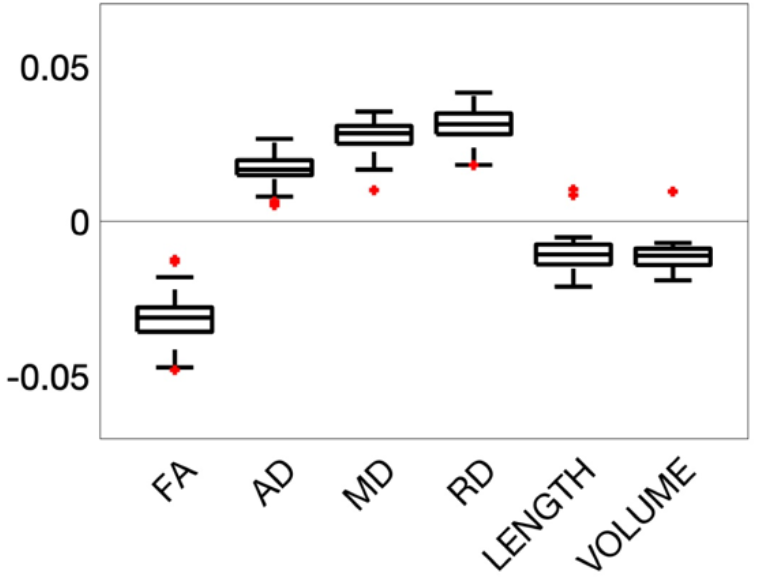
Changes in superficial white matter. The beta coefficient from linear mixed effects modeling across all studied U-fiber pathways is shown in boxplot form (for statistically significant results only). In general, diffusivities show positive associations with age, while FA, length, and volume measures show negative associations with age.

### Visualizing change across superficial white matter

To visualize where changes in SWM occur during aging, all pathways are visualized, colored coded according to percent change per year, and shown in **Figure 6**. Again, SWM pathways throughout the entire cortex show statistically significant increases in diffusivities with age, of ~0.2-0.4% change per year, while FA shows decreases of similar magnitude per year. Notably, microstructural features show greatest changes in frontal and temporal lobes, with minimal changes in pre- and post-central gyri. Changes in length and volume are more sparse, with decreases in length with age observed throughout the entire brain, while decreases in volume with age are denser in the frontal lobe.

**Figure 6.**
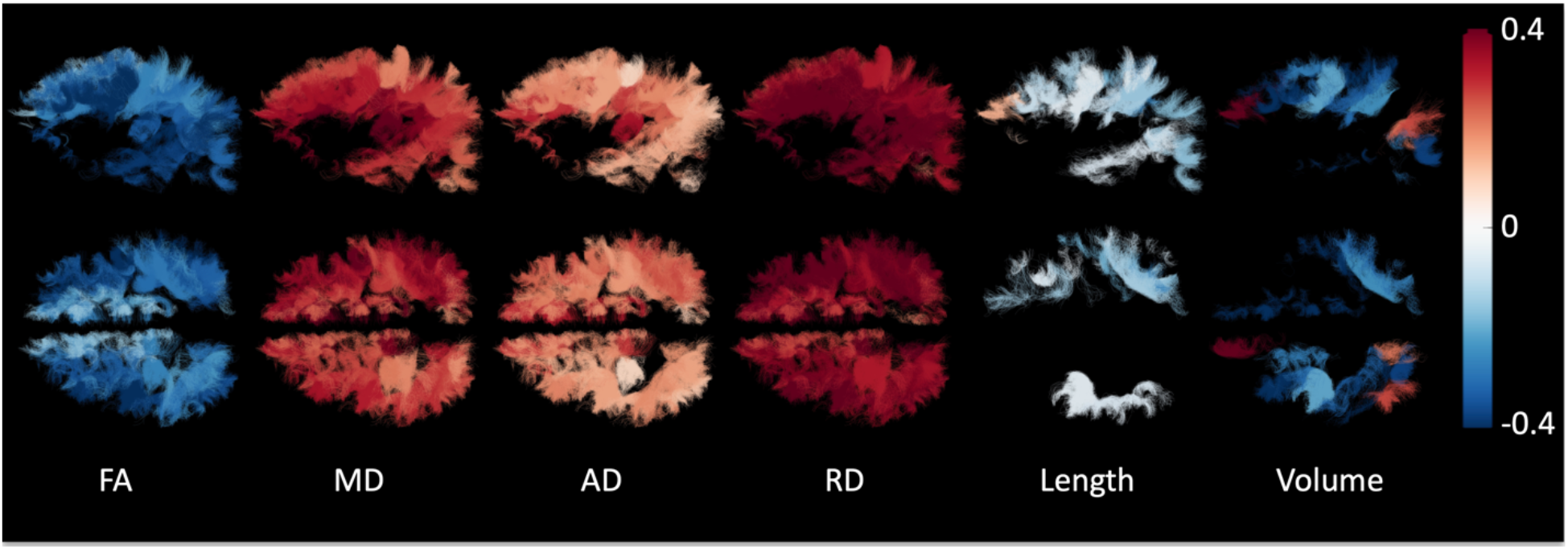
Percent change per year from the population mean shown as color-coded streamlines on an example subject. Bundles are only shown if statistically significant trends with age are observed.

An alternative visualization is shown in **Figure 7**, where each cortical region is color-coded based on the percent-change per year of all SWM fibers connecting that label (note that a single cortical region can be associated with multiple U-fiber systems). Again, clear patterns are observed in SWM associated with frontal and temporal lobes, including larger decreases in FA and increases in all diffusivities. Interestingly, SWM of the pre- and post-central gyri, while indicating less change per year in microstructural features, stand out as the largest decreases in length and volume per year.

**Figure 7.**
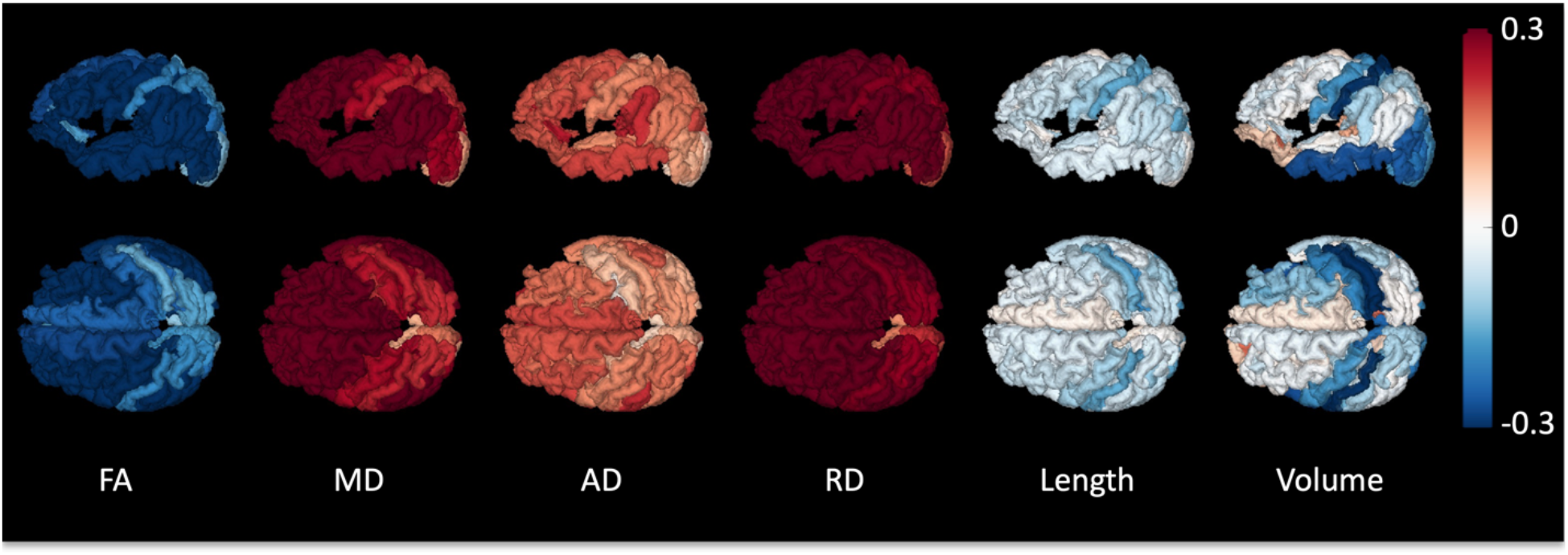
Percent change per year from the population mean for short superficial U-fibers connecting individual regions of interest. Regions of an example subject are color-coded based on the population-averaged percent change per year of all fibers connecting that label.

## Discussion

Here, we have used multiple large, longitudinal and cross-sectional datasets, and innovations in tractography generation and filtering, to characterize U-fiber systems in 3 aging cohorts, describing microstructural features and for the first time, macrostructural features. Our main findings are that (1) diffusivities show positive associations with age, while anisotropy has negative associations with age, in SWM throughout the entire brain, (2) larger microstructural changes were observed in the frontal, temporal, and temporoparietal regions, (3) measures of U-fiber geometry and length decrease with age, and (4) changes in length and volume were more heterogenous, with prominent effects seen at the pre- and post-central gyri.

### Superficial white matter in aging

Compared to the long-range association, projection, and commissural pathways, SWM of the brain has been underexplored in the literature, in both healthy and abnormal aging. Recently, due to advances in software and tools to study SWM, studies of these systems have started to increase. For a thorough review on SWM tractography analysis and applications, see work by Guevara et al. [10]. Of note, there have been few studies of SWM in aging using diffusion MRI. In a study of 141 healthy individuals (18-86 years old), Nazeri et al. [42] found widespread negative relationships of FA with age, in agreement with our results. To do this, they generated a population-based SWM template, and used this to perform a tract-based spatial statistics (TBSS) style analysis. Similarly, in a cohort of 65 individuals (18-74 years old) Phillips et al. [41] found age-related reductions in FA and increases in RD and AD across large areas of SWM, with results more pronounced in the frontal SWM compared to the posterior and ventral brain regions, and they interpreted this as an increased vulnerability to the aging process. Rather than tractography, this was done using white matter/gray matter surface-based alignment from structural MRI data and probing the DTI indices across the population along this boundary. Finally, using tractography and manually placed regions of interest on 69 subjects (22-84 years old), and focusing on prefrontal connections, Malykhin et al. [43] found significant decreases in FA starting at ~60 years of age, in both SWM and association/commissural pathways. The use of tractography also enabled volumetric analysis, where both long range and short-range fiber systems showed decreased volumes with age.

Motivated by these works in SWM, the current study takes advantage of innovations in tractography and U-fiber segmentation, and incorporates multiple large cross-sectional and longitudinal cohorts totaling >1200 participants and >2700 sessions to study SWM throughout the entire brain. Specifically, constrained spherical deconvolution [59], in combination with probabilistic tractography [49] has become prevalent in state-of-the art studies of the human connectome and individual fiber bundles. Combining this with anatomical constraints [60] and subsequent filtering [52] enables robust delineation of white matter systems underneath most of the cortex (**Figure 1**), in alignment with current knowledge of SWM. Similar methodology has been shown to result in reproducible streamlines [12], making studies of clinical cohorts plausible. Further, we include several large datasets on aging, making this the largest cohort to date to study these fibers in any clinical study.

### What changes and where

The observed associations with age include decreased FA, volume, length, and increased axial, radial, and mean diffusivities. The biological mechanism for these age-related changes is not entirely clear, due to the high sensitivity (and low specificity) of these DTI measures to various features of tissue microstructure. In general, these observations in white matter (in both health and disease) have been attributed to various biological mechanisms. Increases in radial and axial diffusivities are often associated with decreased axonal packing [61, 62], allowing for increased diffusivity in all orientations, as well as myelin thinning which may be observed as increased radial diffusivity [63, 64]. The low sensitivity of DTI can potentially be overcome with multi-compartment modeling, which may allow disentangling neurite densities, compartmental changes, and geometrical configurations. For example, a SWM study of individuals with young onset Alzheimer’s disease (using the white matter and gray matter boundary to define regions, as in [41]) found that these individuals exhibited decreased FA and increased diffusivities [65]. However, the use of a multi-compartment tissue model (in this case the neurite orientation dispersion and density imaging model [66], showed both a decreased neurite volume fraction and higher dispersion index, suggesting both a loss of myelinated fibers and greater dispersion (less coherent organization) of these SWM systems. While these studies were able to detect differences in extreme neurodegenerative cases, we found that these systems are sensitive in aging individuals without cognitive impairment as well. Future studies should implement similar modeling, in combination with the tractography generation and segmentation utilized in this study, to improve biological specificity of changes in healthy aging.

Identifying where changes occur during age may facilitate studying the underpinnings of cognitive and motor changes, and aid in identifying networks that are susceptible to disease and disorder. Here, much like previous studies [6, 41, 67–70] in gray matter, white matter pathways, and axonal diameters, there is a clear anterior-to-posterior gradient in changes of microstructure across age. The frontal lobe is comprised of functional networks recruited for a diverse range of cognitive problems, and disruption is associated with age-related declines in cognitive processes [71]. Our study confirms that in addition to gray matter, and the larger white matter pathways, the U-fibers of the frontal lobe also indicate strong age-related trends. future work should investigate relationships between these neuroimaging features and age-related declines in cognition.

### Towards painting a complete picture of brain aging

Noninvasive MR-imaging has slowly led to a convergence of evidence of structural and functional changes in aging. The main findings from decades of research are that the brain shrinks in overall volume and the ventricular system expands in volume [22]. The pattern of changes is heterogenous, as described here and elsewhere [22], with most analyses suggesting a 0.5%-1% reduction in volume per year in most areas of the brain. The changes in volume are related to neuronal loss, neuronal shrinkage, decreased length of myelinated axons in white matter and reduction of synapses in the gray matter. Finally, structural changes in healthy aging mediate, or explain, domain-specific cognitive decline in individuals both with and without cognitive impairment [29, 30]. The results of this study highlight that SWM cannot be ignored when forming a complete picture of brain aging. In addition, variation of these systems across populations may enable subject-specific analysis and identification of atypical structure, which may be used to study subject-specific function.

### Limitations and future direction

Because of the lack of studies on SWM, there are a number of research directions that can benefit from these methodologies. Understanding not only the relationship between SWM and the cortex, but also the SWM and long-range pathways would further our understanding of the complex interactions of the aging brain. Additionally, tractometry [72–74] or high dimensional analysis of the brain, which has been shown to enable single-subject inference [72], may benefit from the additional set of features provided by SWM. Understanding which features of the brain change first is paramount to understanding differences in disease. SWM has found relevant application in cohorts with autism, schizophrenia, and Alzheimer’s disease, [10] and may further benefit from a comprehensive examination of the structural changes of the brain including both white and gray matter geometric analysis and microstructure analysis. Similarly, inclusion of cognitive and motor variables will facilitate linking function to structure. Finally, studies of SWM may help identify challenges for traditional fiber tractography of the long-range fibers – characterizing where these systems occur may facilitate challenges associated with gyral biases [11, 75, 76] and bottlenecks in streamline propagation that lead to creation of false positive pathways [77–80].

Several limitations should be acknowledged. First, while the use of multiple datasets allowed a large sample size, the use of different datasets with different acquisitions is known to result in very different quantitative indices of microstructure and macrostructure [54–58]. However, we included dataset as a variable in our mixed effects models, and consider this an advantage to the current study which shows these effects generalize across all data. Second, the data used is neither high angular resolution nor high spatial resolution, and future studies should utilize higher resolution datasets (e.g., the Human Connectome Project [81]), which may reduce variability in quantification, and enable studies across the entire lifetime. Third, we chose simple linear mixed effects modelling, whereas changes across a lifespan have been shown to be nonlinear – therefore we chose to focus our analysis on age 50+. Fourth, there are several methods to segment and study U-fibers, both with and without tractography [10, 14, 82, 83], and we could have chosen different streamline generation and clustering algorithms. We expect that results will be similar, but not exactly the same, with the use of different methodologies for virtual dissection [84]. Finally, while U-fiber atlases do exist [14, 15, 83, 85, 86], we choose to include all “U-shaped” fiber systems that exist within a certain percent of the studied population. This does not guarantee the existence of true anatomical connections, but has been used in the literature as an indicator of reliability of results.

## Conclusion

Here, we have used a large, longitudinal dataset, and innovations in tractography generation and filtering, to characterize U-fiber systems in an aging cohort, describing microstructural features and for the first time, macrostructural features. We find robust associations with age for all features, across many fiber systems. These features, and their normal variations with age, may be useful for characterizing abnormal aging, and, in combination with larger association pathways and gray matter microstructural features, lead to insight into fundamental mechanisms associated with aging and cognition.

## Statements and Declarations

### Author Contributions

All authors contributed to the study conception and design. Data collection was performed by the Baltimore Longitudinal Study of Aging at the National Institutes of Aging, and the Vanderbilt Memory & Aging Project (VMAP). All authors commented on previous versions of the manuscript. All authors read and approved the final manuscript.

### Data Availability

Derived microstructure and macrostructure features, for all pathways and subjects, along with demographic information, are made available at (link upon acceptance) for VMAP and CAMCAN datasets. Data from the BLSA are available on request from the BLSA website (http://blsa.nih.gov). All requests are reviewed by the BLSA Data Sharing Proposal Review Committee and may also be subject to approval from the NIH institutional review board.

### Ethic Approval

All human datasets from Vanderbilt University were acquired after informed consent under supervision of the appropriate Institutional Review Board. All additional datasets are freely available and unrestricted for non-commercial research purposes. This study accessed only de-identified patient information.

### Consent to participate

Informed consent was obtained from all individual participants included in the study.

### Appendix

Below, we give the abbreviated names used in the manuscript and figure captions, and the freesurfer-based name as given in FreeSurferColorLUT.txt. Here, U-fibers connect one cortical region to another indicated by a “:” in the abbreviation.

| Abbreviation | Freesurfer-based nomenclature |
| --- | --- |
| l g fr mid:l g fr mid | ctx lh G front middle---ctx lh G front middle |
| l g fr sup:l g s cing | ctx lh G front sup---ctx lh G and S cingul-Ant |
| l g fr sup:l g fr mid | ctx lh G front sup---ctx lh G front middle |
| l g fr sup:l g fr sup | ctx lh G front sup---ctx lh G front sup |
| l g prec:l g postc | ctx lh G precentral---ctx lh G postcentral |
| l g prec:l g prec | ctx lh G precentral---ctx lh G precentral |
| l s cent:l g prec | ctx lh S central---ctx lh G precentral |
| l s cent:l s cent | ctx lh S central---ctx lh S central |
| l s fr inf:l g fr mid | ctx lh S front inf---ctx lh G front middle |
| l s fr inf:l s fr inf | ctx lh S front inf---ctx lh S front inf |
| l s fr mid:l g fr mid | ctx lh S front middle---ctx lh G front middle |
| l s fr sup:l g fr mid | ctx lh S front sup---ctx lh G front middle |
| l s fr sup:l g fr sup | ctx lh S front sup---ctx lh G front sup |
| l s fr sup:l s fr sup | ctx lh S front sup---ctx lh S front sup |
| l s orbshape:l s orbshape | ctx lh S orbital-H Shaped---ctx lh S orbital-H Shaped |
| l s pcal:l g fr sup | ctx lh S pericallosal---ctx lh G front sup |
| l s precinpart:l s fr inf | ctx lh S precentral-inf-part---ctx lh S front inf |
| l s precinpart:l s precinpart | ctx lh S precentral-inf-part---ctx lh S precentral-inf-part |
| l s presupport:l g prec | ctx lh S precentral-sup-part---ctx lh G precentral |
| l s presupport:l s presupport | ctx lh S precentral-sup-part---ctx lh S precentral-sup-part |
| r g fr mid:r g fr mid | ctx rh G front middle---ctx rh G front middle |
| r g fr sup:r g s cing | ctx rh G front sup---ctx rh G and S cingul-Ant |
| r g fr sup:r g fr mid | ctx rh G front sup---ctx rh G front middle |
| r g fr sup:r g fr sup | ctx rh G front sup---ctx rh G front sup |
| r g orb:r g orb | ctx rh G orbital---ctx rh G orbital |
| r g prec:r g prec | ctx rh G precentral---ctx rh G precentral |
| r s centr:r g prec | ctx rh S central---ctx rh G precentral |
| r s fr inf:r g fr mid | ctx rh S front inf---ctx rh G front middle |
| r s fr inf:r s fr inf | ctx rh S front inf---ctx rh S front inf |
| r s fr mid:r g fr mid | ctx rh S front middle---ctx rh G front middle |
| r s fr mid:r s fr mid | ctx rh S front middle---ctx rh S front middle |
| r s fr sup:r g fr mid | ctx rh S front sup---ctx rh G front middle |
| r s fr sup:r g fr sup | ctx rh S front sup---ctx rh G front sup |
| r s fr sup:r s fr sup | ctx rh S front sup---ctx rh S front sup |
| r s orbshape:r g orb | ctx rh S orbital-H Shaped---ctx rh G orbital |
| r s pcal:r g fr sup | ctx rh S pericallosal---ctx rh G front sup |
| r s precinpart:r s fr inf | ctx rh S precentral-inf-part---ctx rh S front inf |
| r s precinpart:r s precinpart | ctx rh S precentral-inf-part---ctx rh S precentral-inf-part |
| r s presupport:r g prec | ctx rh S precentral-sup-part---ctx rh G precentral |
| r s presupport:r s presupport | ctx rh S precentral-sup-part---ctx rh S precentral-sup-part |
| l g par infang:l g par infang | ctx lh G pariet inf-Angular---ctx lh G pariet inf-Angular |
| l g par infsup:l g par infsup | ctx lh G pariet inf-Supramar---ctx lh G pariet inf-Supramar |
| l g par sup:l g par sup | ctx lh G parietal sup---ctx lh G parietal sup |
| l g postc:l g postc | ctx lh G postcentral---ctx lh G postcentral |
| l s inpar ptrans:l g par infang | ctx lh S intrapariet and P trans---ctx lh G pariet inf-Angular |
| l s inpar ptrans:l g par sup | ctx lh S intrapariet and P trans---ctx lh G parietal sup |
| l s inpar ptrans:l s inpar ptrans | ctx lh S intrapariet and P trans---ctx lh S intrapariet and P trans |
| l s postc:l g postc | ctx lh S postcentral---ctx lh G postcentral |
| l s postc:l s postc | ctx lh S postcentral---ctx lh S postcentral |
| r g par infang:r g par infang | ctx rh G pariet inf-Angular---ctx rh G pariet inf-Angular |
| r g par infsup:r g par infsup | ctx rh G pariet inf-Supramar---ctx rh G pariet inf-Supramar |
| r g par sup:r g par sup | ctx rh G parietal sup---ctx rh G parietal sup |
| r latfistpost:r g par infsup | ctx rh Lat Fis-post---ctx rh G pariet inf-Supramar |
| r s centr:r s cent | ctx rh S central---ctx rh S central |
| r s inpar ptrans:r g par infang | ctx rh S intrapariet and P trans---ctx rh G pariet inf-Angular |
| r s inpar ptrans:r g par sup | ctx rh S intrapariet and P trans---ctx rh G parietal sup |
| r s inpar ptrans:r s inpar ptrans | ctx rh S intrapariet and P trans---ctx rh S intrapariet and P trans |
| r s paro occ:r s paro occ | ctx rh S parieto occipital---ctx rh S parieto occipital |
| r s postc:r g postc | ctx rh S postcentral---ctx rh G postcentral |
| r s postc:r s inpar ptrans | ctx rh S postcentral---ctx rh S intrapariet and P trans |
| r s postc:r s postc | ctx rh S postcentral---ctx rh S postcentral |
| l g octemp mling:l g octemp mling | ctx lh G oc-temp med-Lingual---ctx lh G oc-temp med-Lingual |
| l s oc mid Lunatus:l g occ mid | ctx lh S oc middle and Lunatus---ctx lh G occipital middle |
| l s oc sup trans:l s oc sup trans | ctx lh S oc sup and transversal---ctx lh S oc sup and transversal |
| l s paro occ:l s paro occ | ctx lh S parieto occipital---ctx lh S parieto occipital |
| r g cuneus:r g cuneus | ctx rh G cuneus---ctx rh G cuneus |
| r g occ mid:r g occ mid | ctx rh G occipital middle---ctx rh G occipital middle |
| r g octemp mling:r g octemp mling | ctx rh G oc-temp med-Lingual---ctx rh G oc-temp med-Lingual |
| r pole occ:r pole occ | ctx rh Pole occipital---ctx rh Pole occipital |
| r s oc mid Lunatus:r g occ mid | ctx rh S oc middle and Lunatus---ctx rh G occipital middle |
| r s oc sup trans:r g occ sup | ctx rh S oc sup and transversal---ctx rh G occipital sup |
| r s oc sup trans:r s oc sup trans | ctx rh S oc sup and transversal---ctx rh S oc sup and transversal |
| l s temp sup:l g par infang | ctx lh S temporal sup---ctx lh G pariet inf-Angular |
| l s temp sup:l g temp mid | ctx lh S temporal sup---ctx lh G temporal middle |
| l s temp sup:l s temp sup | ctx lh S temporal sup---ctx lh S temporal sup |
| r g temp mid:r g temp mid | ctx rh G temporal middle---ctx rh G temporal middle |
| r latfistpost:r latfistpost | ctx rh Lat Fis-post---ctx rh Lat Fis-post |
| r s temp sup:r g par infang | ctx rh S temporal sup---ctx rh G pariet inf-Angular |
| r s temp sup:r g temp mid | ctx rh S temporal sup---ctx rh G temporal middle |
| r s temp sup:r s temp sup | ctx rh S temporal sup---ctx rh S temporal sup |
| l g s cing:l g s cing | ctx lh G and S cingul-Ant---ctx lh G and S cingul-Ant |
| r g s cing:r g s cing | ctx rh G and S cingul-Ant---ctx rh G and S cingul-Ant |

